# Immune cells lacking Y chromosome have widespread dysregulation of autosomal genes

**DOI:** 10.1101/673459

**Authors:** Jan P. Dumanski, Jonatan Halvardson, Hanna Davies, Edyta Rychlicka-Buniowska, Jonas Mattisson, Behrooz Torabi Moghadam, Noemi Nagy, Kazimierz Węglarczyk, Karolina Bukowska-Strakova, Marcus Danielsson, Paweł Olszewski, Arkadiusz Piotrowski, Erin Oerton, Aleksandra Ambicka, Marcin Przewoźnik, Łukasz Bełch, Tomasz Grodzicki, Piotr L. Chłosta, Stefan Imreh, Vilmantas Giedraitis, Lena Kilander, Jessica Nordlund, Adam Ameur, Ulf Gyllensten, Åsa Johansson, Alicja Józkowicz, Maciej Siedlar, Alicja Klich-Rączka, Janusz Jaszczyński, Stefan Enroth, Jarosław Baran, Martin Ingelsson, John R. B. Perry, Janusz Ryś, Lars A. Forsberg

**Affiliations:** Dept. of Immunology, Genetics and Pathology and Science for Life Laboratory, Uppsala University, Uppsala, Sweden; Faculty of Pharmacy and 3P Medicine Laboratory, International Research Agendas Programme, Medical University of Gdańsk, Gdańsk, Poland; International Research Agendas Programme, 3P Medicine Laboratory, Medical University of Gdańsk, Gdańsk, Poland; Dept. of Cell and Molecular Biology, Karolinska Institutet, Stockholm, Sweden; Dept. of Clinical Immunology, Institute of Paediatrics, Jagiellonian University, Collegium Medicum, Kraków, Poland; MRC Epidemiology Unit, School of Clinical Medicine, University of Cambridge, Cambridge, UK; Dept. of Tumour Pathology, Maria Skłodowska-Curie Memorial Cancer Centre and Institute of Oncology, Kraków Branch, Kraków, Poland; Dept. and Clinic of Urology, Jagiellonian University, Collegium Medicum, Kraków, Poland; Dept. and Clinic of Internal Medicine and Gerontology, Jagiellonian University, Collegium Medicum, Kraków, Poland; Dept. Oncology-Pathology, Karolinska Institutet, Stockholm, Sweden; Dept. of Public Health and Caring Sciences/ Geriatrics, Uppsala University, Uppsala, Sweden; Dept. of Medical Sciences, Uppsala University, Uppsala, Sweden; Dept. of Medical Biotechnology, Faculty of Biochemistry, Biophysics and Biotechnology, Jagiellonian University, Kraków, Poland; Dept. of Urology, Maria Skłodowska-Curie Memorial Cancer Centre and Institute of Oncology, Kraków Branch, Kraków, Poland; The Beijer Laboratory, Uppsala University, Uppsala, Sweden

## Abstract

Mosaic loss of chromosome Y (LOY) in leukocytes has been associated with many diseases, yet it remains unclear whether this form of clonal mosaicism exerts a direct physiological effect. Here we perform single-cell and bulk RNA sequencing in leukocytes, observing considerable variation in the rate of LOY across individuals, cell types and disease state. Cells with LOY demonstrated a profound degree of transcriptional dysregulation impacting ∼500 autosomal genes. These genes are preferentially involved in immune functions but also encode proteins with roles in other diverse biological processes. Our findings highlight a surprisingly broad role for chromosome Y challenging the view of it as a “genetic wasteland”. Furthermore, they support the hypothesis that altered immune function in leukocytes is a mechanism directly linking LOY to disease.

## Introduction

For over 50 years it has been noted that chromosome Y is frequently lost in the leukocytes of ageing men,^1,2^ representing the most commonly observed form of clonal mosaicism. Our recent work demonstrated that 1 in 5 men in the UK Biobank study have detectable LOY in blood,^3,4^ reaching a prevalence of 57% in 93 year old men.^5^ Furthermore, LOY is considerably more common in peripheral blood leukocytes vs. other tissues.^5,6^ These observations have been accompanied by epidemiological studies linking LOY in blood to numerous disease outcomes, including all-cause mortality, Alzheimer’s disease, various forms of cancer, autoimmune conditions, age-related macular degeneration, cardiovascular disease, type 2 diabetes and obesity.^6-15^

Recent large-scale population studies have highlighted a heritable component to LOY and begun identifying individual genetic determinants. The largest to date, studying 205,011 men in UK Biobank, identified 156 genetic loci associated with LOY, which were preferentially found near genes involved in cell-cycle regulation, cancer susceptibility, somatic drivers of tumour growth and cancer therapy targets.^4^ The emerging picture is that LOY is in part determined by genetic predisposition to deficiencies in DNA damage response – either through genetic effects that promote chromosome mis-segregation or failure in the molecular machinery to detect and appropriately deal with this damage. The remaining part might be due to other risk factors, e.g. smoking and environmental hazards.^16-18^

There are two main, not mutually exclusive hypotheses that could explain the link between LOY in blood and risk for disease – either loss of a Y chromosome in leukocytes exerts a direct physiological effect, and/or LOY in leukocytes is a barometer of broader genomic instability in other cell types. Evidence for the latter emerged with the observation that genetic susceptibility to LOY influences non-haematological health outcomes in women (who are XX, hence eliminating a direct Y effect). This observation does not however preclude a direct effect of LOY that may explain some of the noted disease associations. Indeed, we recently reported that expression of *TCL1A* was dysregulated in CD19+ B-lymphocytes missing the Y chromosome, providing a proof-of-concept that LOY may not be functionally neutral in these cells.^4^ Our current study substantially expands this observation, demonstrating that LOY leads to profound and widespread dysregulation of gene expression in leukocytes. Furthermore, the biological nature of these dysregulated genes directly supports the hypothesis that altered immune response may be a mechanism linking LOY in immune cells to disease.

## Results

To investigate the functional consequences of LOY, we studied changes in gene expression associated with LOY in vivo and in vitro (see **methods**). The single-cell transcriptome analysis (scRNAseq) of peripheral blood mononuclear cells (PBMCs) collected from 29 aging men generated a pooled dataset encompassing 73,606 cells. The Y chromosome contains 64 protein coding genes: 45 in the male specific region (MSY) and 19 in the pseudo-autosomal regions (PARs). We detected normal expression of 20 protein coding genes on chromosome Y across all white blood cells (**Fig. S1**), enabling the classification of single cells with or without LOY. We identified single cells with LOY in all 29 subjects studied with scRNAseq and observed that the distribution of LOY varied substantially between cell types (**Fig. S2**). Specifically, the frequency of LOY in NK cells, monocytes, B- and T lymphocytes were 27% (range 7-87%), 23% (range 7-87%), 7% (range 2-40%) and 3% (range 1-6%), respectively. We also performed bulk RNA sequencing (RNAseq) on 134 samples of sorted cell fractions (NK cells, monocytes and granulocytes) collected from 51 subjects, including the 29 subjects studied with scRNAseq (**Fig. S2**).

We observed a high concordance in LOY estimates generated from pairwise samples studied in vivo with scRNAseq, RNAseq and DNA based technologies (**Fig. S3**). Furthermore, we studied in vitro changes of gene expression using RNAseq in 13 lymphoblastoid cell lines (LCLs) with or without LOY (**Figs. S4 and S5**). Concordant levels of LOY were observed in LCLs studied with SNP-array as well as with a new method for LOY analysis^19^ using droplet digital PCR targeting a 6 bp sequence difference between the *AMELY* and *AMELX* genes (Pearson’s correlation coefficient = 0.9961; details not shown).

### Many autosomal genes are dysregulated in LOY cells

To identify <u>L</u>OY <u>A</u>ssociated <u>T</u>ranscriptional <u>E</u>ffects (LATE) we tested for differential gene expression between the LOY and non-LOY cellular populations (**Table 1, Fig. 1, Fig. S6**, see **methods**). As expected, MSY gene expression was greatly decreased in the bulk RNAseq on sorted cells and absent in the single cell data (as this was the LOY definition for single cells). A corresponding analysis of the genes located in the PARs also showed a decrease in transcript abundance with increasing levels of LOY. However, the decrease in PAR genes was not as distinct as for the MSY genes, which was expected because of sustained expression of the chromosome X-copy (**Fig. 1, Fig. S4**). Outside of the Y chromosome, we found evidence for 489 autosomal LATE genes and 10 on the non-PAR X chromosome across the two expression datasets and all types of leukocytes, from in vivo studied samples (**Fig. 1, Table 1, Table S1**). Of these, 75 were identified in the single cell data and 420 in the bulk RNAseq and we observed overexpression as well as underexpression of specific autosomal LATE genes. The autosomal genes showing the largest LATE were *LYPD2* and *IL1R2*, displaying 8.6 higher (DE = 3.11, FDR = 0.0011) and 2.5 lower (DE = −1.30, FDR = 0.0612) abundance of transcripts, respectively. A considerable number of genes were independently identified as showing LATE in different cell types and by both technologies (**Table S2**). Co-expression analysis showed that autosomal genes that are normally co-expressed with MSY genes displayed a higher level of differential expression in single cells with LOY compared with control genes (**Fig. S4**, Wilcoxon rank sum test: *p*=0.0021, see **methods**). Results from in vitro studied LCL’s further support that LOY is associated with altered autosomal gene expression using a complementary approach (**Figs. S4 and S5**, Kolmogorov–Smirnov test: D=1.0, *p*=0.0016).

**Table 1.** Number of autosomal genes with LOY associated transcriptional effects (LATE) observed in samples studied in vivo and in vitro using two RNA sequencing methods.

| Cell type | Results from RNAseq analysis * |  | Results from scRNAseq analysis * |  |
| --- | --- | --- | --- | --- |
|  | N. of expressed genes in samples without LOY | N. of LATE genes in samples with LOY | N. of expressed genes in normal non-LOY cells | N. of LATE genes in single cells with LOY |
| **NK cells** | 9974 | 298 | 3021 | 37 |
| **Monocytes** | 9891 | 80 | 3634 | 28 |
| **Granulocytes** | 7497 | 57 | - | - |
| **T lymphocytes** | - | - | 2798 | 13 |
| **B lymphocytes** | - | - | 3112 | 3 |
| **LCLs** | 9549 | 757 | - | - |
| **Number of genes #** | 11472 | 1155 | 4631 | 75 |
| **Number of LATE genes in vivo #** |  | 420 |  | 75 |
| **Total number of autosomal LATE genes detected in vivo by RNAseq and/or scRNAseq # = 489** |  |  |  |  |

\* Results from RNAseq and scRNAseq analyses are shown separately for each cell type. The detected number (N.) of expressed genes as well as the number of LATE genes detected in different cell types using FDR<0.1 are shown.

**Fig. 1.**
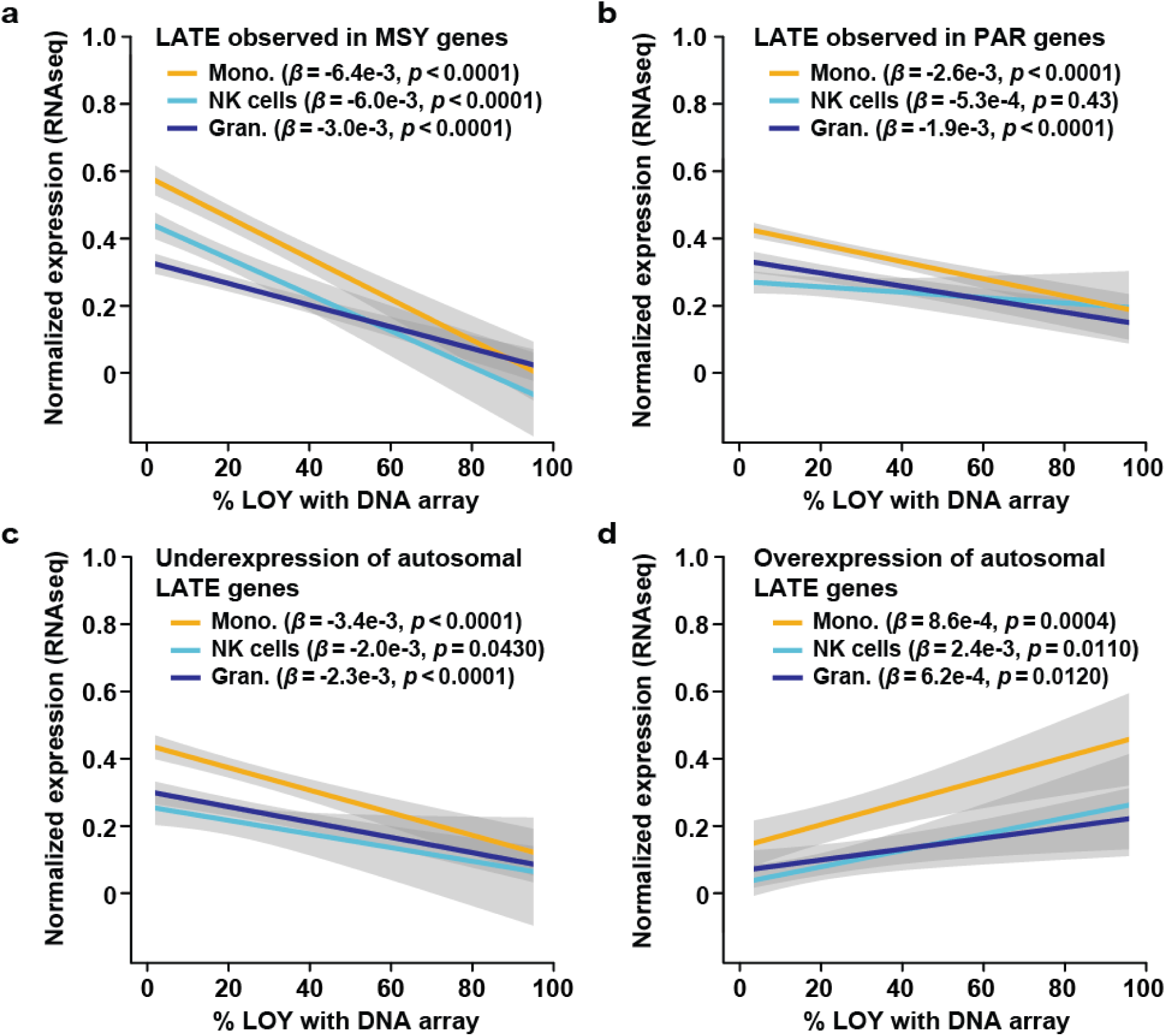
LOY Associated Transcriptional Effect (LATE) in leukocytes in vivo. On the X-axes are the level of LOY mosaicism (estimated from DNA by SNP-array from three sorted cell populations) and on the Y-axes are the normalized level of gene expression estimated from the same samples using RNAseq. Panel a display the average expression of 6 genes located in the male-specific part of chromosome Y (MSY) as a function of LOY. Panel b show corresponding analysis of 13 genes located in the pseudo-autosomal regions of chromosomes X and Y (PAR). Panels c and d illustrate the finding of autosomal LATE genes, i.e. genes located on other chromosomes showing reduced or increased abundance of transcripts in samples with LOY. Panel c display the average expression of the ten most underexpressed autosomal LATE genes and the ten most overexpressed autosomal LATE genes is shown in panel d. Grey areas represent the standard error of linear regression models and beta (*β*) with confidence estimate (*p*) is shown. Abbreviations: Mono. = monocytes, Gran. = granulocytes.

In the monocytes and NK-cells that were studied by two RNA sequencing methods, the identified LATE genes showed an overall concordance in the direction of change in gene expression between technologies (binomial sign test; monocytes: *p*=0.030, NK-cells: *p*=0.001). The level and direction of differential expression of LATE genes detected independently by both technologies are illustrated in **Fig. 2**. Among these, two genes were detected in both cell types and by both methods, i.e. upregulation of the *LY6E* gene located on chromosome 8 and downregulation of the PAR gene *CD99*.

**Fig. 2.**
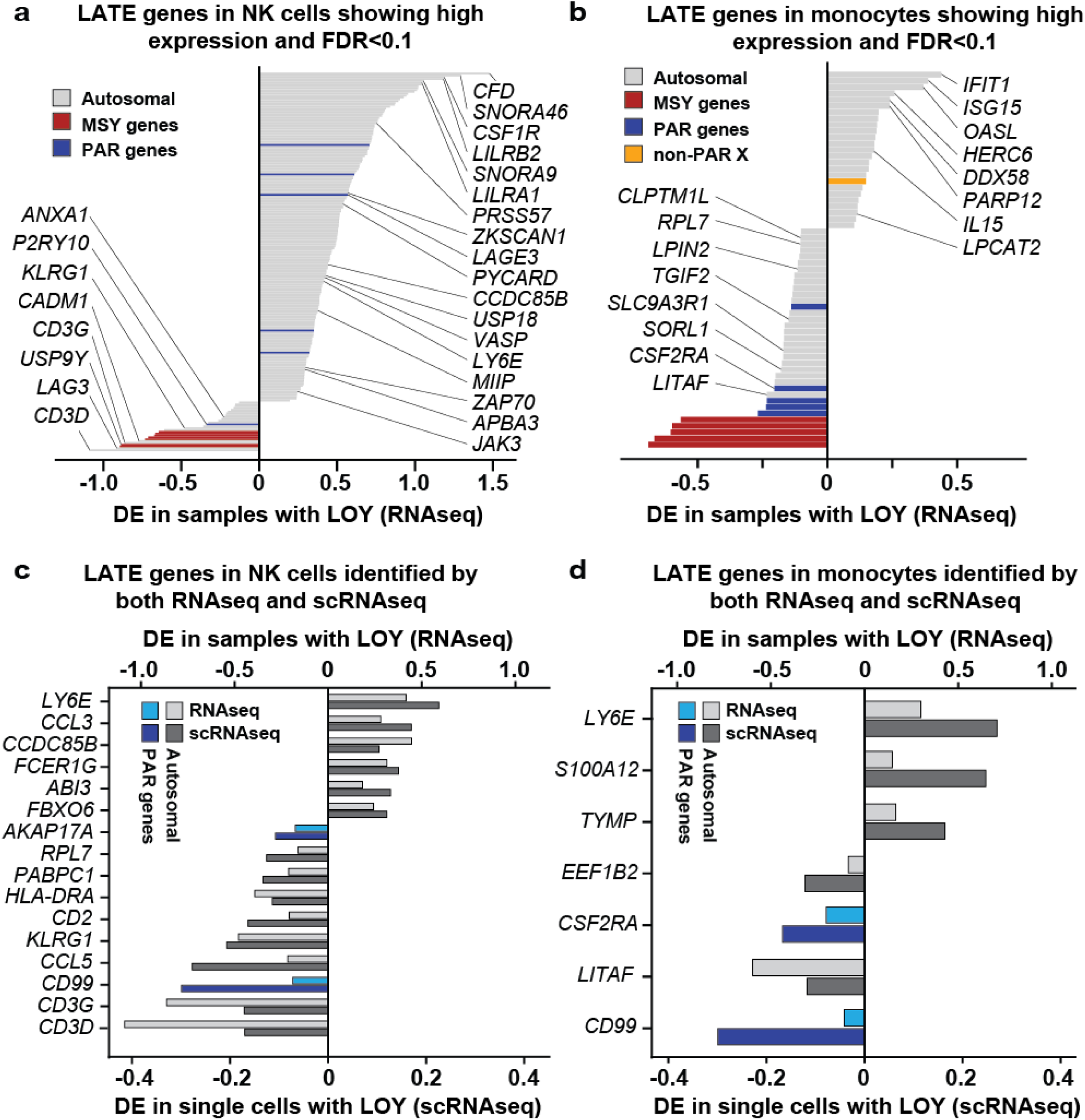
Differential expression (DE) of specific genes as an effect of LOY in NK cells and monocytes, studied by both RNAseq and scRNAseq. Panels a and b illustrate the level of DE in 206 and 60 autosomal LATE genes identified in each cell type after correction for multiple testing (FDR<0.1) and with at least an average of 100 reads per gene in samples without LOY. Names are shown for LATE genes known to be linked with immune system functions and/or cancer and/or development of Alzheimer’s disease. Panels c and d display DE observed in autosomal and PAR genes that was identified as LATE genes by both RNAseq and scRNAseq at a 0.05 α-level. MSY genes were excluded because of lack of expression in cells without chromosome Y. Independent identification of LATE was observed for 16 and 7 genes in the NK cells and monocytes, respectively. The autosomal gene *LY6E* and the PAR gene *CD99* displayed LATE in both cell types and by both technologies.

Furthermore, we observed that the fraction of LATE genes within specific cell types was substantially larger than the fraction of LATE genes shared between different subsets of cells, in both the RNAseq dataset (ANOVA: *F*_*1,8*_ = 95.5, *p* < 0.0001) and in the scRNAseq dataset (ANOVA: *F*_*1,8*_ = 13.7, *p* = 0.0061). For instance, up to 15% of normally expressed genes showed LATE in NK cells, but only about 2% of LATE genes were shared between NK cells and granulocytes or monocytes (**Fig. S7**). This result emphasize that transcriptomic alterations associated with LOY varies between different types of immune cells, likely because each cell type normally express a set of genes that is only partially overlapping with the gene expression of other cell types.

### Biological pathways implicated by identified LATE genes

The 489 autosomal LATE genes had twice as many (*p*<1×10^−16^) biological connections amongst each other than expected by chance for a matched randomly selected set of genes. Furthermore, LATE genes were not randomly distributed throughout the genome, with clusters of genes evident (**Figure S8**). This was most pronounced in the 11q13.1 region, where we identified 17 LATE genes (13 over-expressed in LOY) in an 8 Mb genomic window (**Table S1**). To explore the potential biological relevance of the identified genes we next performed pathway enrichment analyses, controlling for the background set of all genes expressed in our tested cell types. A total of 306 Gene Ontology and Reactome terms/pathways (FDR < 0.05) were significantly over-represented, the majority of which focused on aspects of RNA processing, immune function and the viral life cycle (**Table S3**). A summary of our literature review for detected LATE genes is presented in **Table S4**.

We next tested whether LATE genes were preferentially located near genetic variants associated with LOY.^4^ Gene set enrichment analysis implemented in MAGENTA demonstrated a modest excess of such associations (1.17 fold enrichment, *p*=0.010), with 41 LATE genes mapping within 300 kb of a genome-wide significant SNP association in 25 genomic regions. In several instances, the proximal LATE gene was the same as that prioritised through gene mapping approaches in the GWAS study. For example, LATE gene cyclin-dependent kinase inhibitor 1C (*CDKN1C*) was linked via expression changes to a nearby LOY-associated variant (rs60808706). Here, the LOY risk increasing allele increases expression of *CDKN1C* and the expression of *CDKN1C* is increased in cells with LOY. As with our previously reported example of TCL1A,^4^ this observation raises the possibility of a bi-directional relationship between LOY and *CDKN1C -* genetically increased expression of the gene promotes LOY and LOY subsequently further dysregulates gene expression in a cell-specific manner, possibly leading to a negative spiral of increased clonal mosaicism via perturbed cell cycle processes.

### LOY in sorted leukocytes from patients with prostate cancer and Alzheimer’s disease

Finally, we investigated the distribution of LOY in six types of sorted leukocytes - CD19+ B lymphocytes, CD4+ T lymphocytes, CD8+ T-cytotoxic lymphocytes, Natural Killer (NK) cells, granulocytes and monocytes - among men diagnosed with Alzheimer’s disease (N=121) or prostate cancer (N=107) vs. healthy controls (N=156). Overall, there was a general trend for LOY to be increased in patients vs. controls, and this rate varied considerably between cell types (**Fig. 3**). Men with Alzheimer’s disease had significantly higher levels of LOY only in CD16+/CD56+ NK cells compared to controls (*p* = 0.0071). The fraction of Alzheimer’s disease cases with high levels of LOY in NK cells was about four times larger compared with controls (not shown) and supports previous observations that LOY is associated with incident Alzheimer’s disease.^9^ Furthermore, the involvement of NK cells in the pathogenesis of Alzheimer’s disease has been suggested before in a range of experimental study designs.^20-28^ In prostate cancer patients, NK cells did not show significantly higher level of LOY. However, in the latter cohort, the CD4+ lymphocytes and granulocytes were more frequently affected by LOY (*p* = 0.033 and *p* = 0.031, respectively). The observation that prostate cancer and Alzheimer’s disease patients were primarily affected with LOY in different types of immune cells supports a disease-specific link. This suggests that associations between LOY and disease cannot solely be attributed to a confounding age-dependent process of genomic instability, which would be expected to equally affect different types of leukocytes. Our results are thus compatible with both hypotheses presented in the introduction regarding aetiology of LOY and disease; accumulation of LOY in leukocytes as a barometer of genomic instability in somatic cells, as well as increasing risk for different types of disease when specific types of immune cells are affected with LOY. Furthermore, we found that patients frequently had LOY in more than one immune cell type. Subjects with LOY in >20% of cells in at least two types of sorted leukocytes was considered LOY-oligoclonal (**Fig. 3** and **Fig. S9**). Overall, the frequency of LOY-oligoclonality was higher in men diagnosed with AD or PC compared with controls (OR=2.74, *p*=0.0026). In the AD cohort, LOY-oligoclonality was significantly higher in patients (OR=2.64, *p*=0.0127) and a difference in the same direction was observed in the PC cohort.

**Fig. 3.**
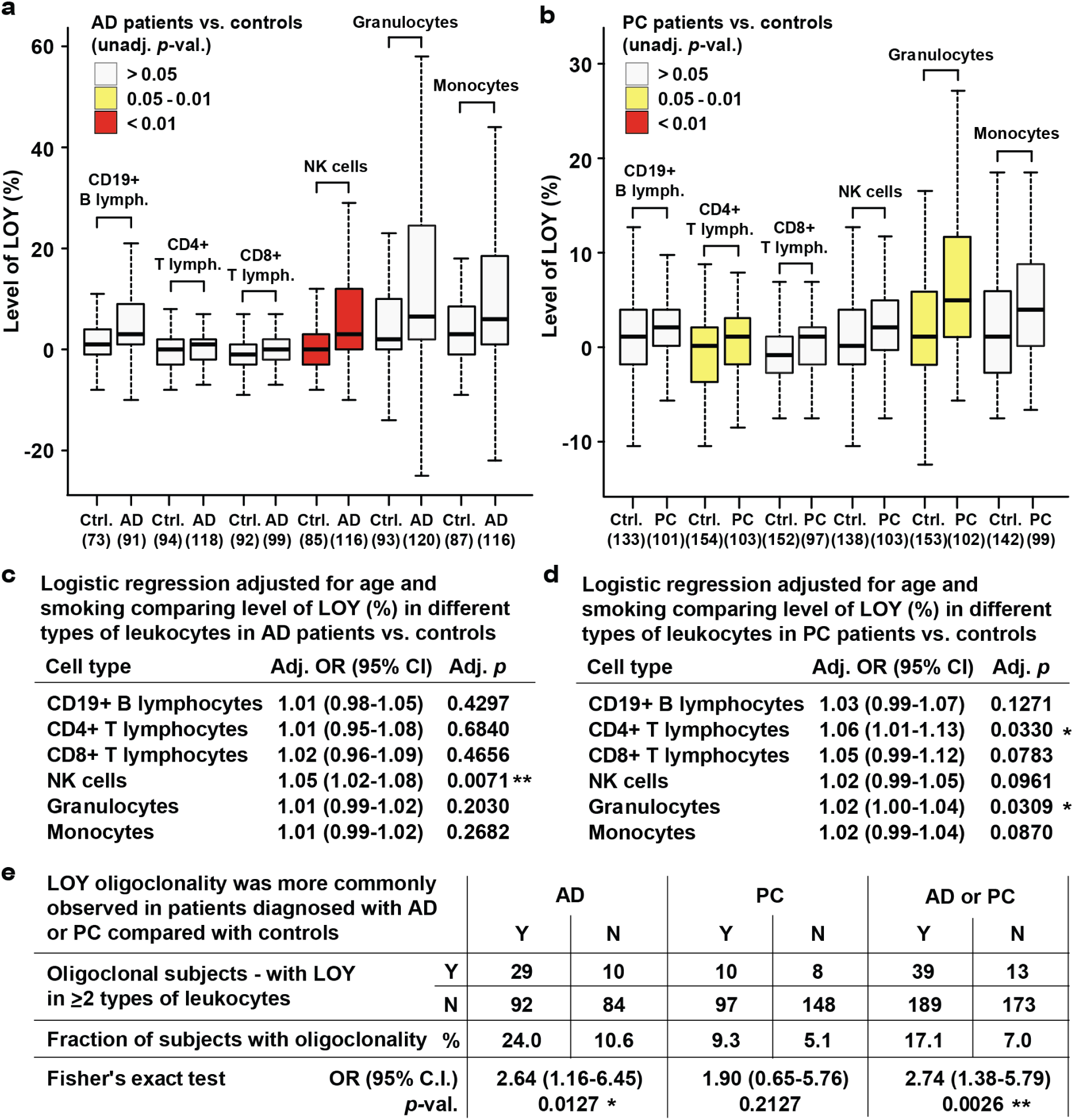
Distribution of LOY in six populations of leukocytes in men diagnosed with Alzheimer’s disease or prostate cancer vs. controls. Leukocytes were sorted by FACS, followed by SNP-array genotyping for each cell fraction and calculation of the percentage of LOY. The numbers in parentheses under the X-axes in panels a and b denote the number of studied subjects for each cell type. Panels a and b as well as panels c and d show results from unadjusted analyses and logistic regression models adjusted for age and smoking, respectively. Panel e shows investigation of LOY oligoclonality in men with AD and PC diagnoses vs. controls. Abbreviations: Ctrl. = control, AD = Alzheimer’s disease, PC = prostate cancer, LR = logistic regression, OR = odds ratio, lymph.= lymphocytes and NK = natural killer.

## Discussion

Our study demonstrates that gene expression is substantially dysregulated in leukocytes with LOY, providing two major insights. Firstly, it highlights a surprisingly broad molecular role for the Y chromosome, which is often considered a “genetic wasteland” with limited involvement in the aetiology of complex traits and diseases. The expression of *SRY* on the Y chromosome is required in mammals to override the development of the default female sex. The mammalian sex chromosomes evolved from a pair of autosomes over the past 300 million years and the Y chromosome has been remarkably stable over the past 25 million years^29,30^. Characterization of human chromosome Y has however been limited by the challenges of sequencing the highly repetitive regions as well as assaying its sequence variation for testing in large population studies. The observation that expression levels of ∼500 autosomal genes change with LOY status suggests that this chromosome may have a broader relevance to regulation of transcriptome and disease than is currently appreciated.^31^ Furthermore, we report that LOY occurs with different frequencies in six sorted fractions of leukocytes in patients with prostate cancer and AD, suggesting there might be a cellular specificity of LOY and supporting a disease-specific link. This cellular specificity of LOY and observation that the fraction of LATE genes within specific cell types was substantially larger than the fraction of LATE genes shared between different subsets of leukocytes (**Fig. S7**) suggests that LOY might have pleiotropic effects. Pleiotropy is defined as an effect of one feature at the genetic level to multiple characteristics at the phenotypic level.^32^ Considering the large number of epidemiological associations between LOY in blood and various disease outcomes, the LOY-pleiotropy might be a plausible concept.

Secondly, our study highlights altered immune cell function that could conceivably link LOY in immune cells directly to disease mechanism(s). The combined number of LATE genes derived from analyses of various cells is surprisingly large, which suggests that a loss of ∼2% of the male haploid genome (via LOY) has a profound impact on cellular homeostasis. Enrichment analyses showed that LATE genes are important for a range of physiological functions (**Table S3)** including many normal functions of the immune system. One of the LATE genes that showed the strongest downregulation was the autosomal gene *LAG3* (**Fig. 2**). The LAG3 protein is a cell surface molecule that functions as an immune checkpoint receptor by binding its main ligand MHC class II with higher affinity than CD4. Cellular proliferation and immune cell activation is regulated by a balance between CD4/LAG3, in a similar fashion to the CTLA4 and PD1 immune checkpoints, where *LAG3* expression suppresses cell activity.^33,34^ The observed low expression of *LAG3* in LOY cells might disrupt the CD4/LAG3 balance by releasing one of the breaks of immune cell activity. Furthermore, it is noteworthy that the well-known immune checkpoint PD-1 signaling pathway is one of the top hits in the enrichment analysis (**Table S3).**

Furthermore, the genes *LY6E* and *CD99* showed LATE by both RNAseq and scRNAseq in NK cells and monocytes (**Fig. 2**). The protein product of *CD99* is a cell surface glycoprotein involved in processes such as leukocyte migration, cell adhesion and apoptosis; e.g. by functioning as a diapedesis-mediating receptor central for migration of monocytes through endothelial junctions.^35,36^ A lower expression of *CD99* in leukocytes with LOY might therefore mitigate extravasation and thus impair the recruitment and movement of leukocytes from circulation towards somatic tissues and sites of disease. Furthermore, the *LY6E* gene was upregulated as an effect of LOY and it has the potential to inhibit inflammatory cytokines and disrupt inflammatory cascades. A survey of more than 130 published clinical studies found that increased expression of *LY6E* is associated with poor survival outcome in multiple malignancies^37^ and it has also been found to be important for drug resistance and tumor immune escape in breast cancer.^38^ Among all autosomal LATE genes, the *IL1R2* gene showed the strongest downregulation in immune cells with LOY. This cytokine receptor is central for orchestrating inflammatory and immune responses by binding of ligands such as interleukin-1α (IL1A), interleukin-1β (IL1B), and interleukin 1 receptor antagonist (IL1Ra), preventing them from binding to other receptors.^39^ Hence, a lower abundance of *IL1R2* in LOY cells could reduce the inhibiting effect of this receptor.

Additionally, several of the LATE genes we identified had diverse and well-described roles in other biological processes, notably *GABRR1, PRLR* and *LEP. GABRR1* encodes a receptor for Gamma Aminobutyric Acid (GABA), the major inhibitory neurotransmitter in the brain. A recent study found these receptors were functionally expressed in human and mouse hematopoietic stem cells and megakaryocyte progenitors.^40^ Overexpression of *GABRR1* in these cells led to increased rates of megakaryocyte and platelet differentiation. In contrast, we observed 1.26 fold (*p*=7.9×10^−5^) increased expression of the receptor in NK cells lacking the Y chromosome, extending the biological role of this receptor in immune cell function. Sex hormone receptors have also been described to have functional effects on haematopoietic stem/progenitor cells^41^ including the prolactin receptor (*PRLR*), which we see over-expressed in monocytes with LOY (**Table S2**). We also observed over-expression of the hunger inhibiting hormone Leptin (LEP) in granulocytes with LOY, consistent with studies describing its lesser known roles in regulating innate immune response.^42^

As mentioned in the introduction, LOY is the most common post-zygotic mutation from analyses of bulk DNA from peripheral blood. The current study adds new information on this as all 29 men (median age 80 years, range 64-94 years) studied by scRNAseq carried leukocytes with LOY. Moreover, as we show here, LOY occurs frequently in blood as oligo-clonal expansions and this could be best explained by independent mutations occurring in progenitors for different lineages of hematopoietic cells. Thus, LOY is more common than so far appreciated and the future analyses of single-cells or sorted-cells from various tissues will be instrumental for establishment of true frequency of this mutation as well as for identification of cell subtypes most often associated with disease. Lastly, we provide results showing LATE of ∼500 autosomal genes, preferentially involved in immune functions but also with roles in other biological processes. In summary, our study provides unique insights into the molecular role of the Y chromosome and how its mosaic loss in leukocytes may directly influence cells functions and impact risk for disease.

## Acknowledgements

We express gratitude to all patients and healthy donors of blood samples from Poland and Sweden. We acknowledge Käthe Ström, Brygida Mazgaj, Bożena Lackowska and Niklas Dahl for help with collection of blood samples from patients and controls as well as Katarina Tegnér and Amanda Raine for assistance with scRNAseq library preparations. We thank Darek Kędra for critical review of the manuscript. This work was supported by grants from the Hjärnfonden, Swedish Cancer Society, the Swedish Research Council, Alzheimerfonden, Konung Gustav V:s och Drottning Viktorias Frimurarestiftelse, the Science for Life Laboratory - Uppsala and the Foundation for Polish Science under the International Research Agendas Programme to J.P.D. as well as from the European Research Council ERC Starting Grant (#679744), the Swedish Research Council (#2017-03762), and Kjell och Märta Beijers Stiftelse to L.A.F. Sequencing was performed by the SNP&SEQ and UGC Technology Platforms at Uppsala University. These facilities are part of the National Genomics Infrastructure (NGI) Sweden and Science for Life Laboratory. The SNP&SEQ Platform is also supported by the Swedish Research Council and the Knut and Alice Wallenberg Foundation.

## Author Contributions

J.P.D. and L.A.F. designed the study and obtained the funding. J.P.D., J.H., J.M., B.T.-M., N.N., M.D., E.O., J.N., Ad.A., S.E., J.R.B.P. and L.A.F. analyzed the data. H.D., E.R.-B., N.N., K.W., K.B.-S., P.O., A.P., Al.A., M.P., L.B., T.G, P.L.C., S.I., V.G., L.K., U.G., Å.J., A.J., M.S., A.K.-R., J.J., S.E., J.B., M.I. and J.R. contributed to data and sample collection. J.P.D., J.H., H.D., E.R.-B., J.M., M.D., S.E., J.R.B.P. and L.A.F. wrote the first draft of the paper. All authors contributed to the final version of the paper.

## REFERENCES

1. Jacobs, P.A., Brunton, M., Court Brown, W.M., Doll, R. & Goldstein, H. Change of human chromosome count distribution with age: evidence for a sex differences. Nature 197, 1080–1 (1963).

2. Pierre, R.V. & Hoagland, H.C. Age-associated aneuploidy: loss of Y chromosome from human bone marrow cells with aging. Cancer 30, 889–94 (1972).

3. Wright, D.J. et al. Genetic variants associated with mosaic Y chromosome loss highlight cell cycle genes and overlap with cancer susceptibility. Nat Genet 49, 674–679 (2017).

4. Thompson, D.J. et al. Genetic predisposition to mosaic Y chromosome loss in blood. Nature 575, 652–657 (2019).

5. Forsberg, L. et al. Mosaic loss of chromosome Y (LOY) in leukocytes matters. Nature Genetics 51, 4–7 (2019).

6. Haitjema, S. et al. Loss of Y Chromosome in Blood Is Associated With Major Cardiovascular Events During Follow-Up in Men After Carotid Endarterectomy. Circ Cardiovasc Genet 10, e001544 (2017).

7. Forsberg, L.A. et al. Mosaic loss of chromosome Y in peripheral blood is associated with shorter survival and higher risk of cancer. Nat Genet 46, 624–8 (2014).

8. Ganster, C. et al. New data shed light on Y-loss-related pathogenesis in myelodysplastic syndromes. Genes Chromosomes Cancer 54, 717–24 (2015).

9. Dumanski, J.P. et al. Mosaic Loss of Chromosome Y in Blood Is Associated with Alzheimer Disease. Am J Hum Genet 98, 1208–19 (2016).

10. Noveski, P. et al. Loss of Y Chromosome in Peripheral Blood of Colorectal and Prostate Cancer Patients. PLoS One 11, e0146264 (2016).

11. Loftfield, E. et al. Predictors of mosaic chromosome Y loss and associations with mortality in the UK Biobank. Sci Rep 8, 12316 (2018).

12. Loftfield, E. et al. Mosaic Y Loss Is Moderately Associated with Solid Tumor Risk. Cancer Res 79, 461–466 (2019).

13. Persani, L. et al. Increased loss of the Y chromosome in peripheral blood cells in male patients with autoimmune thyroiditis. J Autoimmun 38, J193–6 (2012).

14. Lleo, A. et al. Y chromosome loss in male patients with primary biliary cirrhosis. J Autoimmun 41, 87–91 (2013).

15. Grassmann, F. et al. Y chromosome mosaicism is associated with age-related macular degeneration. Eur J Hum Genet 27, 36–41 (2019).

16. Dumanski, J.P. et al. Smoking is associated with mosaic loss of chromosome Y. Science 347, 81–83 (2015).

17. Wong, J.Y.Y. et al. Outdoor air pollution and mosaic loss of chromosome Y in older men from the Cardiovascular Health Study. Environ Int 116, 239–247 (2018).

18. Liu, Y. et al. Polycyclic aromatic hydrocarbons exposure and their joint effects with age, smoking, and TCL1A variants on mosaic loss of chromosome Y among coke-oven workers. Environ Pollut, Published online, doi: 10.1016/j.envpol.2019.113655 (2019).

19. Danielsson, M. et al. Longitudinal changes in the frequency of mosaic chromosome Y loss in peripheral blood cells of aging men varies profoundly between individuals. Eur J Hum Genet, Published online, doi: 10.1038/s41431-019-0533-z (2019).

20. Masera, R.G. et al. Mental deterioration correlates with response of natural killer (NK) cell activity to physiological modifiers in patients with short history of Alzheimer’s disease. Psychoneuroendocrinology 27, 447–61 (2002).

21. Solerte, S.B., Cravello, L., Ferrari, E. & Fioravanti, M. Overproduction of IFN-gamma and TNF-alpha from natural killer (NK) cells is associated with abnormal NK reactivity and cognitive derangement in Alzheimer’s disease. Ann N Y Acad Sci 917, 331–40 (2000).

22. Schindowski, K. et al. Apoptosis of CD4+ T and natural killer cells in Alzheimer’s disease. Pharmacopsychiatry 39, 220–8 (2006).

23. Jadidi-Niaragh, F., Shegarfi, H., Naddafi, F. & Mirshafiey, A. The role of natural killer cells in Alzheimer’s disease. Scand J Immunol 76, 451–6 (2012).

24. Araga, S., Kagimoto, H., Funamoto, K. & Takahashi, K. Reduced natural killer cell activity in patients with dementia of the Alzheimer type. Acta Neurol Scand 84, 259–63 (1991).

25. Le Page, A. et al. NK Cells are Activated in Amnestic Mild Cognitive Impairment but not in Mild Alzheimer’s Disease Patients. J Alzheimers Dis 46, 93–107 (2015).

26. Le Page, A. et al. Role of the peripheral innate immune system in the development of Alzheimer’s disease. Exp Gerontol 107, 59–66 (2018).

27. Mrdjen, D. et al. High-Dimensional Single-Cell Mapping of Central Nervous System Immune Cells Reveals Distinct Myeloid Subsets in Health, Aging, and Disease. Immunity 48, 380–395 e6 (2018).

28. Korin, B. et al. High-dimensional, single-cell characterization of the brain’s immune compartment. Nat Neurosci 20, 1300–1309 (2017).

29. Cortez, D. et al. Origins and functional evolution of Y chromosomes across mammals. Nature 508, 488–93 (2014).

30. Bellott, D.W. et al. Mammalian Y chromosomes retain widely expressed dosage-sensitive regulators. Nature 508, 494–9 (2014).

31. Case, L.K. et al. The Y chromosome as a regulatory element shaping immune cell transcriptomes and susceptibility to autoimmune disease. Genome Res 23, 1474–85 (2013).

32. Paaby, A.B. & Rockman, M.V. The many faces of pleiotropy. Trends Genet 29, 66–73 (2013).

33. Huang, C.T. et al. Role of LAG-3 in regulatory T cells. Immunity 21, 503–13 (2004).

34. Workman, C.J. et al. Lymphocyte activation gene-3 (CD223) regulates the size of the expanding T cell population following antigen activation in vivo. J Immunol 172, 5450–5 (2004).

35. Schenkel, A.R., Mamdouh, Z., Chen, X., Liebman, R.M. & Muller, W.A. CD99 plays a major role in the migration of monocytes through endothelial junctions. Nat Immunol 3, 143–50 (2002).

36. Vestweber, D. How leukocytes cross the vascular endothelium. Nat Rev Immunol 15, 692–704 (2015).

37. Luo, L. et al. Distinct lymphocyte antigens 6 (Ly6) family members Ly6D, Ly6E, Ly6K and Ly6H drive tumorigenesis and clinical outcome. Oncotarget 7, 11165–93 (2016).

38. AlHossiny, M. et al. Ly6E/K Signaling to TGFbeta Promotes Breast Cancer Progression, Immune Escape, and Drug Resistance. Cancer Res 76, 3376–86 (2016).

39. Wang, D. et al. Structural insights into the assembly and activation of IL-1beta with its receptors. Nat Immunol 11, 905–11 (2010).

40. Zhu, F. et al. The GABA receptor GABRR1 is expressed on and functional in hematopoietic stem cells and megakaryocyte progenitors. Proc Natl Acad Sci U S A 116, 18416–18422 (2019).

41. Abdelbaset-Ismail, A. et al. Human haematopoietic stem/progenitor cells express several functional sex hormone receptors. J Cell Mol Med 20, 134–46 (2016).

42. Procaccini, C. et al. Leptin as immune mediator: Interaction between neuroendocrine and immune system. Dev Comp Immunol 66, 120–129 (2017).

43. Love, M.I., Huber, W. & Anders, S. Moderated estimation of fold change and dispersion for RNA-seq data with DESeq2. Genome Biol 15, 550 (2014).

44. Butler, A., Hoffman, P., Smibert, P., Papalexi, E. & Satija, R. Integrating single-cell transcriptomic data across different conditions, technologies, and species. Nat Biotechnol 36, 411–420 (2018).

45. Szklarczyk, D. et al. STRING v11: protein-protein association networks with increased coverage, supporting functional discovery in genome-wide experimental datasets. Nucleic Acids Res 47, D607–D613 (2019).

46. Turner, S.D. qqman: an R package for visualizing GWAS results using Q-Q and manhattan plots. Published online, doi: doi:10.1101/005165 (2014).

47. Segre, A.V. et al. Common inherited variation in mitochondrial genes is not enriched for associations with type 2 diabetes or related glycemic traits. PLoS Genet 6(2010).

48. Carlson, M. org.Hs.eg.db: Genome wide annotation for Human. (2019).

49. Fresno, C. & Fernandez, E.A. RDAVIDWebService: a versatile R interface to DAVID. Bioinformatics 29, 2810–1 (2013).

50. Mi, H., Muruganujan, A., Ebert, D., Huang, X. & Thomas, P.D. PANTHER version 14: more genomes, a new PANTHER GO-slim and improvements in enrichment analysis tools. Nucleic Acids Res 47, D419–D426 (2019).

