## supplemental file for "Immune cells lacking Y chromosome have widespread dysregulation of autosomal genes"

***Item*** ***Page(s)***

Fig. S1………………….... 2

Fig. S2………………….... 3

Fig. S3………………….... 4

Fig. S4………………….... 5-6

Fig. S5……………...…..... 7

Fig. S6………………….... 8-9

Fig. S7………………….... 10

Fig. S8……………...…..... 11-12

Fig. S9……………...…..... 13-15

Tables S1-S4 are available online.

**Fig. S1**

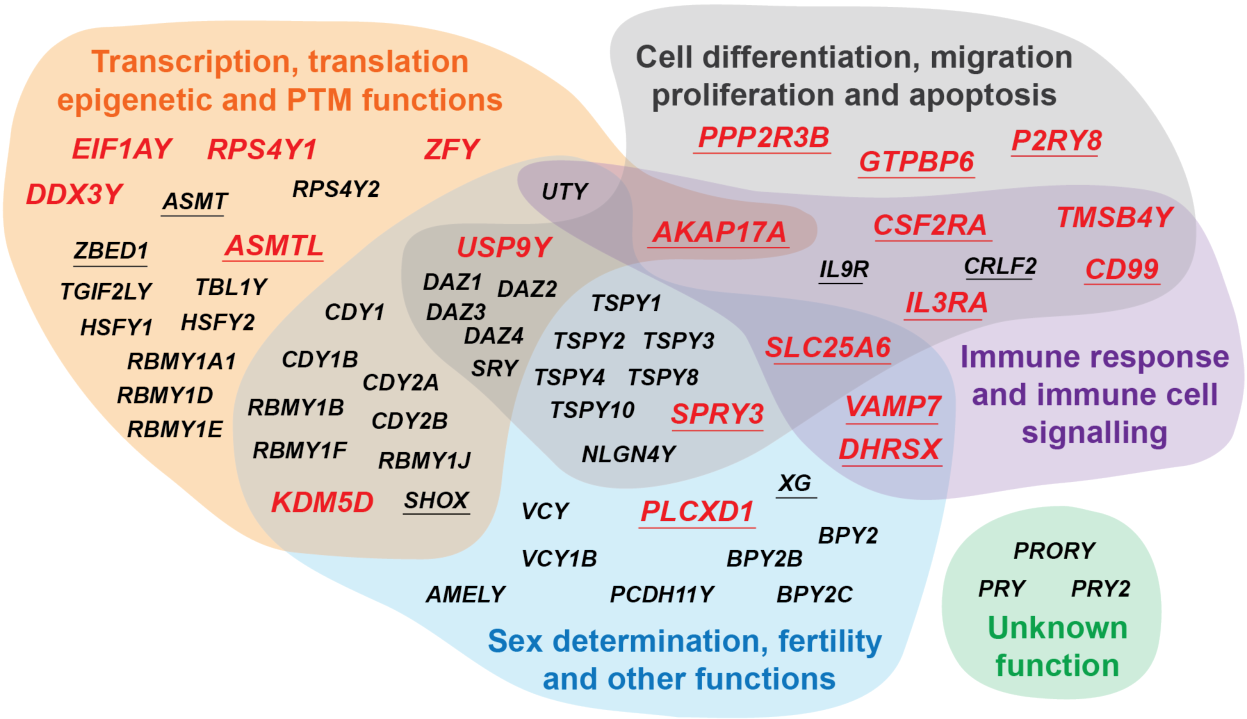

**Supplementary Fig. 1.** Summary of genes from chromosome Y that are expressed in leukocytes. Normal functions of the 64 protein coding genes on chromosome Y, 19 located in the pseudo-autosomal regions (PAR, underlined) and 45 in the male specific region of chromosome Y (MSY). Red color indicates 20 genes expressed in leukocytes studied here using RNAseq, 13 in PARs and 7 in MSY regions. Gene ontology (GO) analyses were performed for each chromosome Y gene to identify known biological functions and the identified GO terms were annotated into four functional categories. The categories were: "Transcription, translation epigenetic and post-translational modifications (PTM) functions" (orange area), "Cell differentiation, migration proliferation and apoptosis" (grey area), "Sex determination, fertility and other functions" (blue area) and "Immune response and immune cell signaling.

Fig. S2

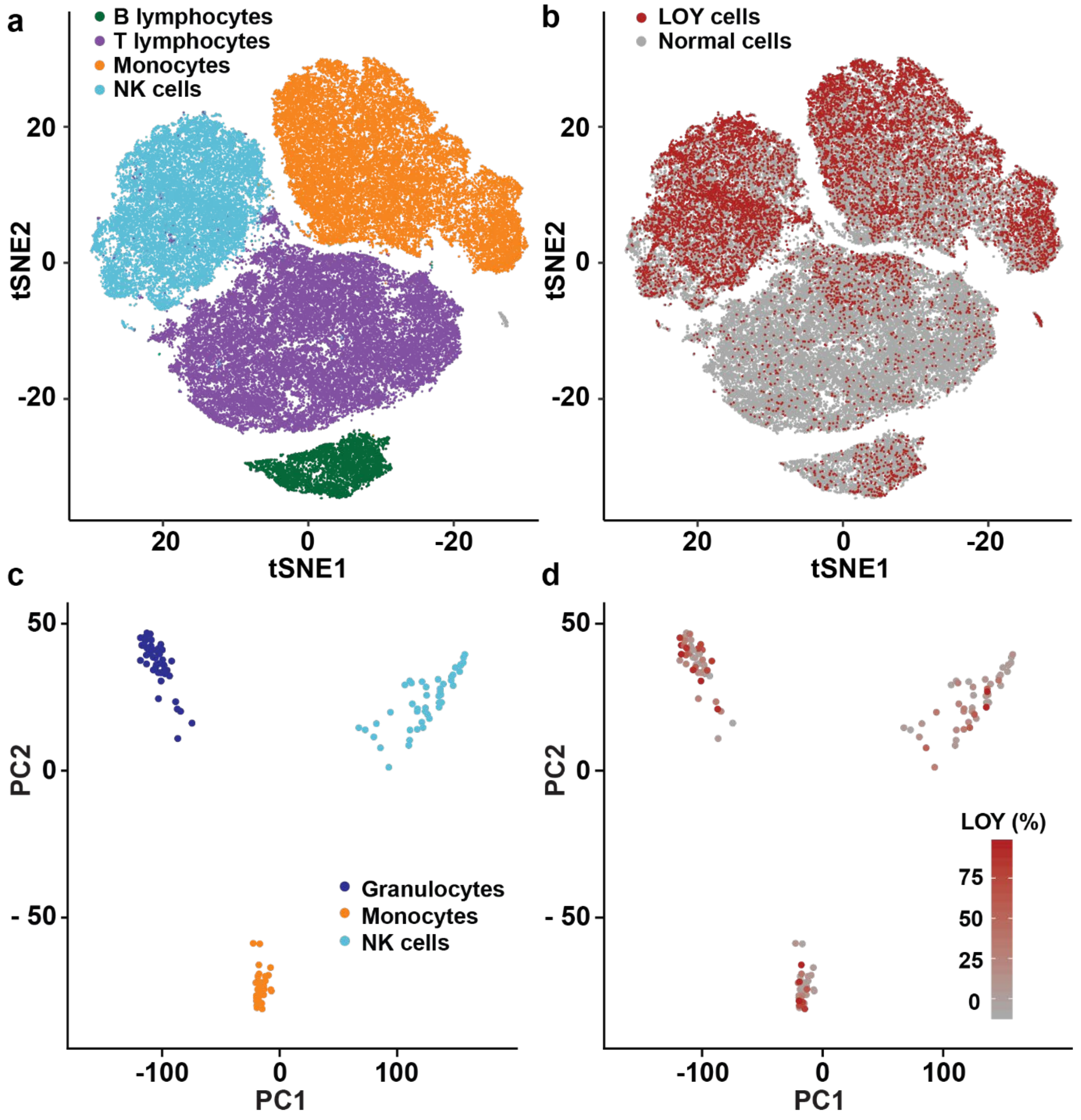

**Supplementary Fig. 2.** Distribution and level of loss of chromosome Y (LOY) in leukocytes from aging men using two RNA-sequencing platforms. Panels a and b show results from single cell RNA sequencing of peripheral blood mononuclear cells (PBMCs) collected from 29 men, 26 diagnosed with Alzheimer's disease. The tSNE plot in panel a shows pooled data from 73,606 PBMCs, each dot representing a single cell, in four cell types that are distinguished by colors. Panel b shows the distribution of cells displaying LOY in different cell types by coloring LOY cells in red and normal cells in grey. Panels c and d show results from bulk RNA sequencing (RNAseq) performed in three cell types sorted by FACS from 51 individuals. Panel c displays a principal component analysis based on global gene expression where each dot represents one cell type in one individual. The distance between dots indicate similarity in gene expression. Panel d shows the level of LOY mosaicism in each sample by a gradient of red and grey where red indicates a high level of LOY.

**Fig. S3**

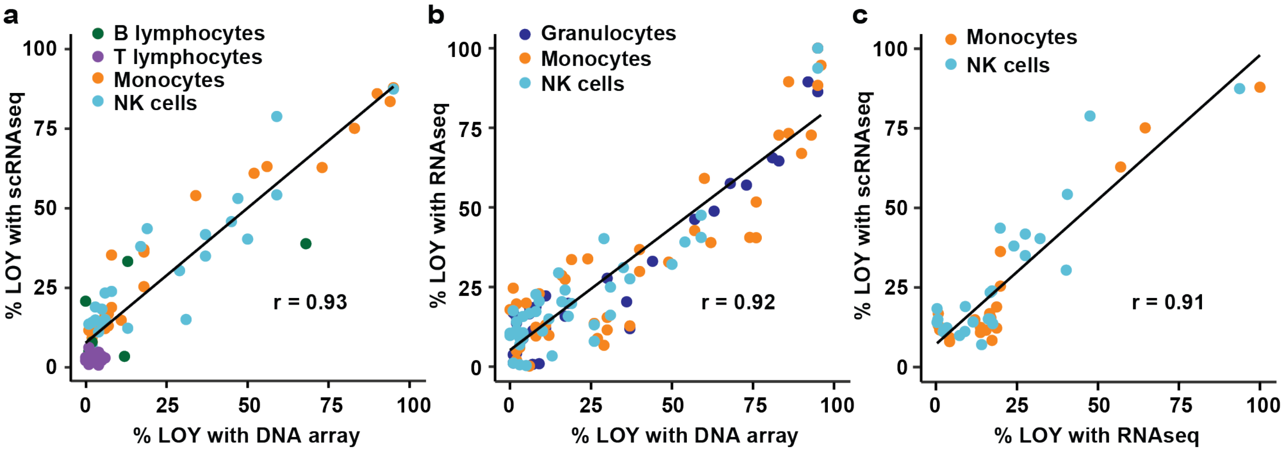

**Supplementary Fig. 3.** Comparison of performance of two RNA-sequencing platforms for estimation of the level of LOY in leukocytes. From the same set of blood samples, LOY analysis was performed in pairwise comparisons between scRNAseq vs. array-based SNP genotyping (panel a), RNAseq vs. array genotyping (panel b) and scRNAseq vs. RNAseq (panel c). For the single cell data, pseudo-bulk samples were created for these comparisons by pooling the single cell data from each cell type for each individual. The Pearson's correlation coefficient for the pairwise analyses were 0.93, 0.92 and 0.91, respectively. Abbreviations: scRNAseq = single cell RNA sequencing, RNAseq = bulk RNA sequencing, r = Pearson's correlation coefficient.

Fig. S4

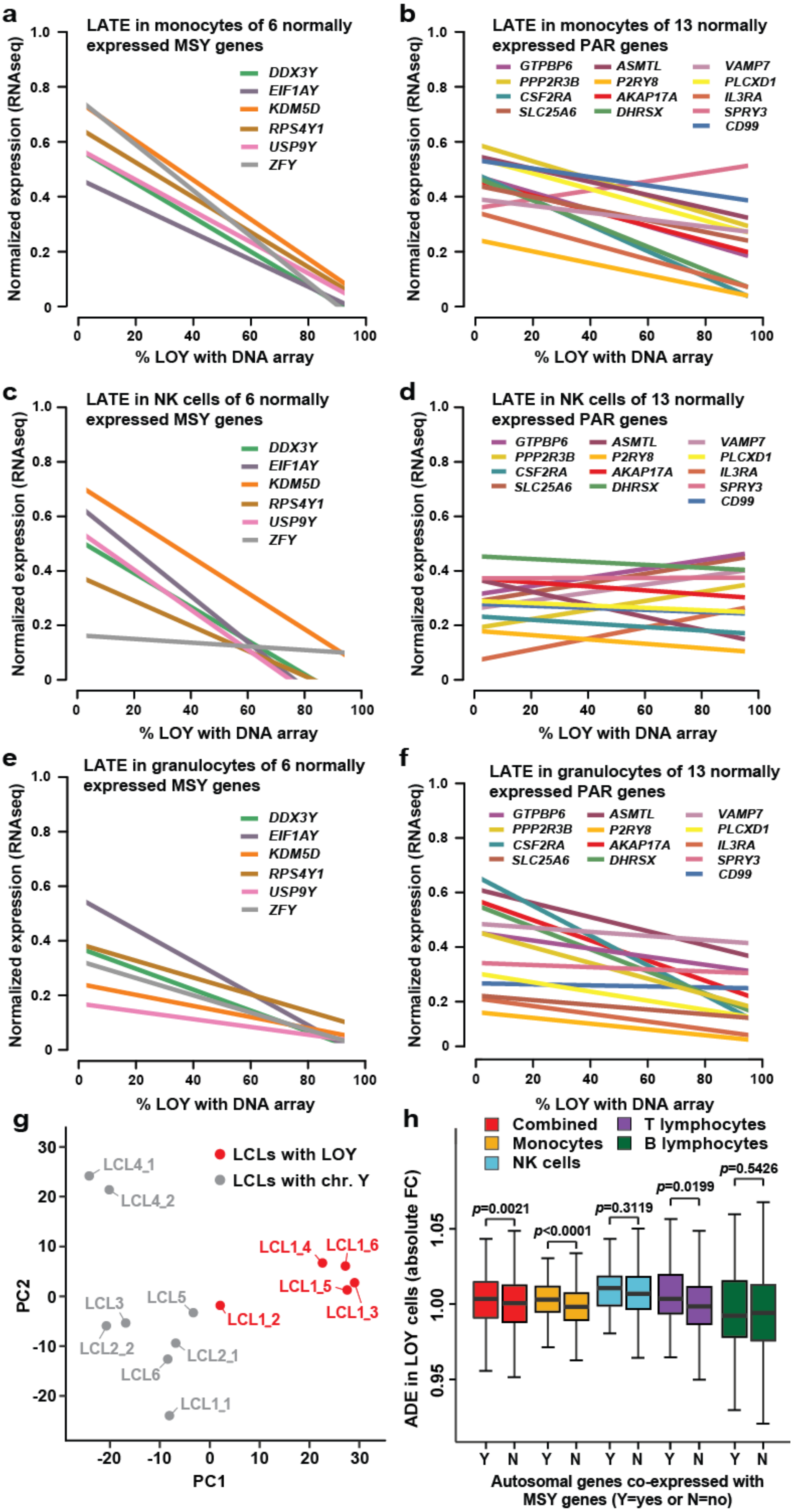
a
LATE in monocytes of 6 normally expressed MSY genes
DDX3Y
EIF1AY
KDM5D
RPS4Y1
USP9Y
ZFY
Normalized expression (RNAseq)
% LOY with DNA array
b
LATE in monocytes of 13 normally expressed PAR genes
GTPBP6
ASMTL
VAMP7
PPP2R3B
P2RY8
PLCXD1
CSF2RA
AKAP17A
IL3RA
SLC25A6
DHRSX
SPRY3
CD99
c
LATE in NK cells of 6 normally expressed MSY genes
d
LATE in NK cells of 13 normally expressed PAR genes
e
LATE in granulocytes of 6 normally expressed MSY genes
f
LATE in granulocytes of 13 normally expressed PAR genes
g
LCLs with LOY
LCLs with chr. Y
LCL4_1
LCL4_2
LCL1_4
LCL1_6
LCL1_5
LCL1_3
LCL1_2
LCL3
LCL5
LCL2_2
LCL2_1
LCL6
LCL1_1
PC1
PC2
h
Combined
T lymphocytes
Monocytes
B lymphocytes
NK cells
p=0.0021
p<0.0001
p=0.3119
p=0.0199
p=0.5426
ADE in LOY cells (absolute FC)
Autosomal genes co-expressed with MSY genes (Y=yes or N=no)

**Supplementary Fig. 4** LOY associated Transcriptional Effect (LATE) *in vivo* and *in vitro*. Fluorescence activated cell sorting was used to isolate monocytes, NK cells and granulocytes from 51 males and in total 47, 46 and 47 samples of each cell type were analyzed, respectively. For each sample, the gene expression was estimated using RNAseq and plotted on the Y-axes and the level of LOY plotted on the X-axes were determined with DNA arrays. In panels a-f the normalized gene expression data for each studied gene are scaled between 0 and 1 to allow comparison of expression levels. Panels a, c and e show that the average expression of the six genes located in the male specific part of chromosome Y (MSY) and normally expressed in all cell types studied (i.e. *RPS4Y1, ZFY, USP9Y, DDX3Y, KDM5D* and *EIF1AY*) was decreasing with increasing level of LOY. The gene *TMSB4Y* was not expressed in all cell types and was therefore not included. Panels b, d and f shows a less pronounced but similar decrease in analyses of 13 genes normally expressed in lymphocytes and located in the pseudo-autosomal regions (PARs) of chromosomes X and Y (i.e. *GTPBP6, PPP2R3B, CSF2RA, SLC25A6, ASMTL, P2RY8, AKAP17A, DHRSX, CD99, VAMP7, PLCXD1, IL3RA* and *SPRY3*). Panel g shows a principal component analysis plot of global gene expression in lymphoblastoid cell lines (LCLs). Red and grey dots represent LCLs with and without LOY, respectively, and donor identity are specified by numbers (i.e. LCL2_1 and LCL2_2 are from donor 2). Distance between dots indicates similarity in global gene expression and analysis of the variance in PC1 shows that LCLs with LOY have altered global autosomal expression compared to LCLs with the Y chromosome intact (Kolmogorov–Smirnov test: D=1.0, $p$=0.0016). Panel h shows results from co-expression analysis and autosomal differential expression (ADE) in single cells with LOY. Autosomal genes that are normally co-expressed with MSY genes displayed a higher level of differential expression in single cells with LOY compared with the control genes without normal co-expression with MSY genes (Wilcoxon rank sum test: $p$=0.0021).

### Fig. S5

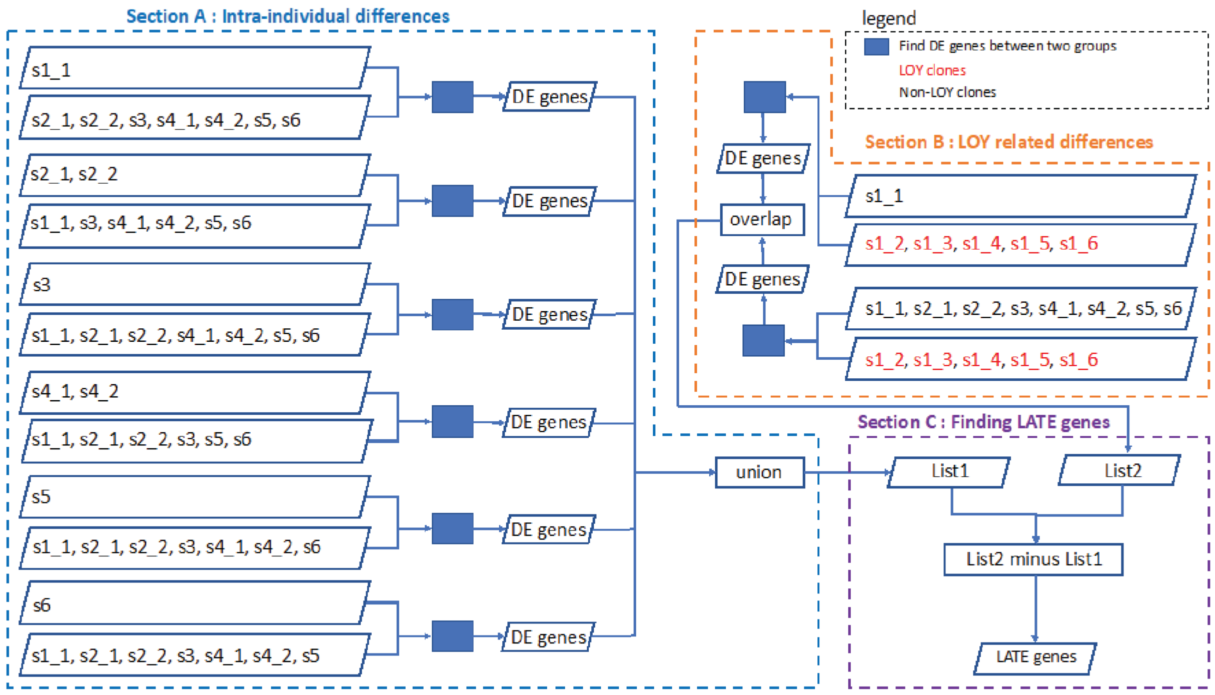

**Supplementary Fig. 5**. The flowchart illustrating analysis pipeline for finding LOY Associated Transcriptional Effect (LATE) genes in lymphoblastoid cell lines (LCLs). LCLs containing 100% LOY cells and no LOY cells are shown in red and black, respectively. Different cell lines derived from the same subject are shown with the same subject ID and an additional number; for instance "s2_1" and "s2_2" are two different cell lines derived from subject s2. The differentially expressed (DE) genes have been called using edgeR by comparing two groups of samples; this process is represented by blue boxes in the chart. In Section A, we compare each non-LOY subject (all cell lines, if there are multiple from the same subject) against all other non-LOY cell lines. The aim is to find the DE genes where the differential expression levels are due to the differences between individual subjects, and not the LOY status. The union of all DE gene lists from Section A generates List1, which is composed of 4175 genes. In Section B, we aim at finding DE genes related to LOY status by comparing: i) all LOY cell lines with all non-LOY clones; and ii) comparing five LOY clones of subject s1 with non-LOY clone from the same subject. The overlap of DE gene lists resulting from (i) and (ii) generates List2 (1531 genes). Finally, in Section C, the list of 904 LATE genes is made by taking the DE genes that exist in List2 but do not exist in List1. The rationale was to eliminate intra-individual differences in expression levels, from LOY-related variation.

### Fig. S6

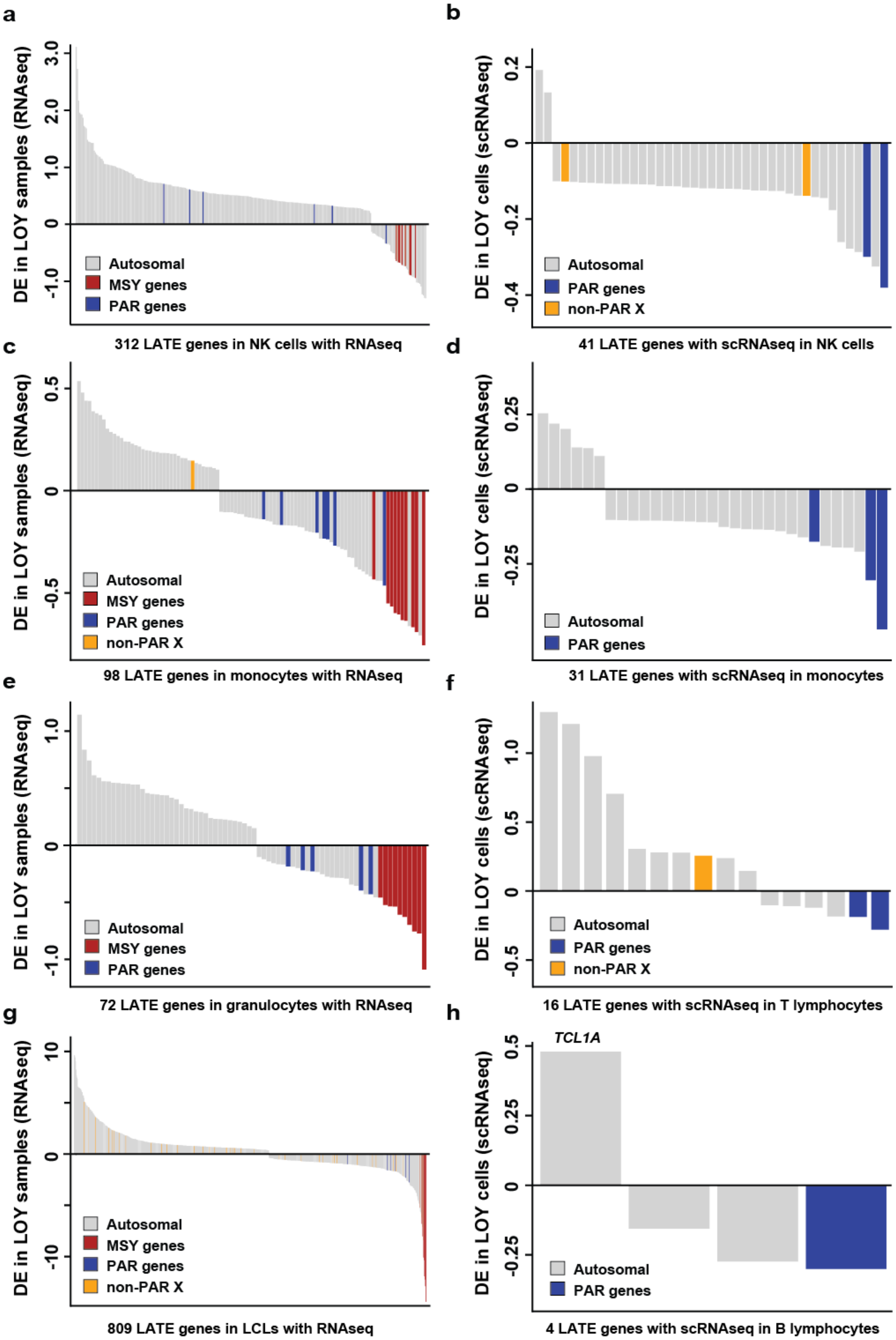

a
DE in LOY samples (RNAseq)
3.0
2.0
1.0
0
-1.0
Autosomal
MSY genes
PAR genes
312 LATE genes in NK cells with RNAseq
b
DE in LOY cells (scRNAseq)
0.2
0
-0.2
-0.4
Autosomal
PAR genes
non-PAR X
41 LATE genes with scRNAseq in NK cells
c
DE in LOY samples (RNAseq)
0.5
0
-0.5
Autosomal
MSY genes
PAR genes
non-PAR X
98 LATE genes in monocytes with RNAseq
d
DE in LOY cells (scRNAseq)
0.25
0
-0.25
Autosomal
PAR genes
31 LATE genes with scRNAseq in monocytes
e
DE in LOY samples (RNAseq)
1.0
0
-1.0
Autosomal
MSY genes
PAR genes
72 LATE genes in granulocytes with RNAseq
f
DE in LOY cells (scRNAseq)
1.0
0.5
0
-0.5
Autosomal
PAR genes
non-PAR X
16 LATE genes with scRNAseq in T lymphocytes
g
DE in LOY samples (RNAseq)
10
0
-10
Autosomal
MSY genes
PAR genes
non-PAR X
809 LATE genes in LCLs with RNAseq
h
TCL1A
DE in LOY cells (scRNAseq)
0.5
0.25
0
-0.25
Autosomal
PAR genes
4 LATE genes with scRNAseq in B lymphocytes

**Supplementary Fig. 6**. Level of differential expression (DE) of LATE genes detected in bulk RNAseq and scRNAseq data. All genes passing correction for multiple testing (FDR<0.1) are shown, i.e. autosomal as well as genes located on the sex chromosomes. Panels a and b, as well as panels c and d, show data from NK cells and monocytes, respectively. Panels e, f, g and h display LATE genes from analysis of granulocytes, T-lymphocytes, LCLs and B-lymphocytes. Genes located in the MSY are indicated in red, PAR genes are indicated in blue, chromosome X genes outside the PAR regions shown in yellow and grey bars show level of differential expression for autosomal genes.

Fig. S7

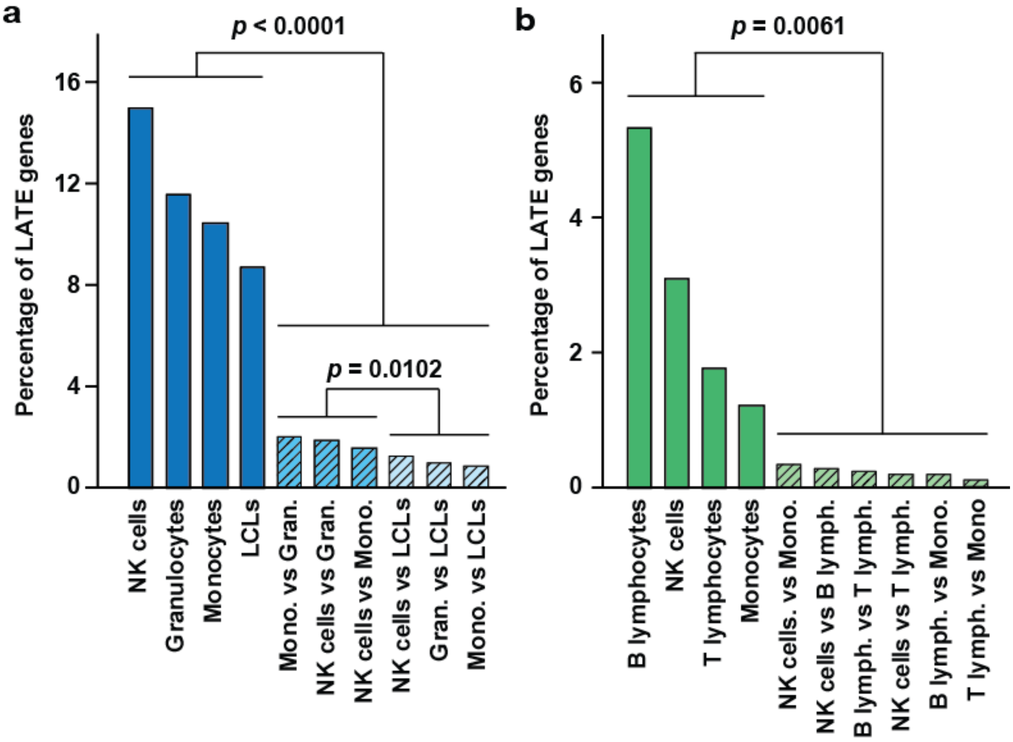

**Supplementary Fig. 7**. The percentage of LATE genes within and between cell types and technologies. A larger fraction of LATE genes within cell types was observed compared with LATE genes shared between cell types. In both datasets, a 0.05 α-level was applied for identification of LATE genes for these analyses. The fraction of genes showing LATE within cell types (open bars) as well as the fraction of LATE genes shared between different cell types (striped bars) was quantified in samples analyzed using RNAseq (panel a) and scRNAseq (panel b). The fraction of shared LATE genes was significantly larger within specific cell types compared with the fraction of LATE genes shared between different types of leukocytes in the RNAseq dataset (ANOVA: $F_{1,8} = 95.5$, $p < 0.0001$) and in the scRNAseq dataset (ANOVA: $F_{1,8} = 13.7$, $p = 0.0061$). The fraction of shared LATE genes was larger also within the *in vivo* collected samples compared with *in vitro* studied LCLs (ANOVA: $F_{1,4} = 20.1$, $p = 0.0102$).

Fig. S8

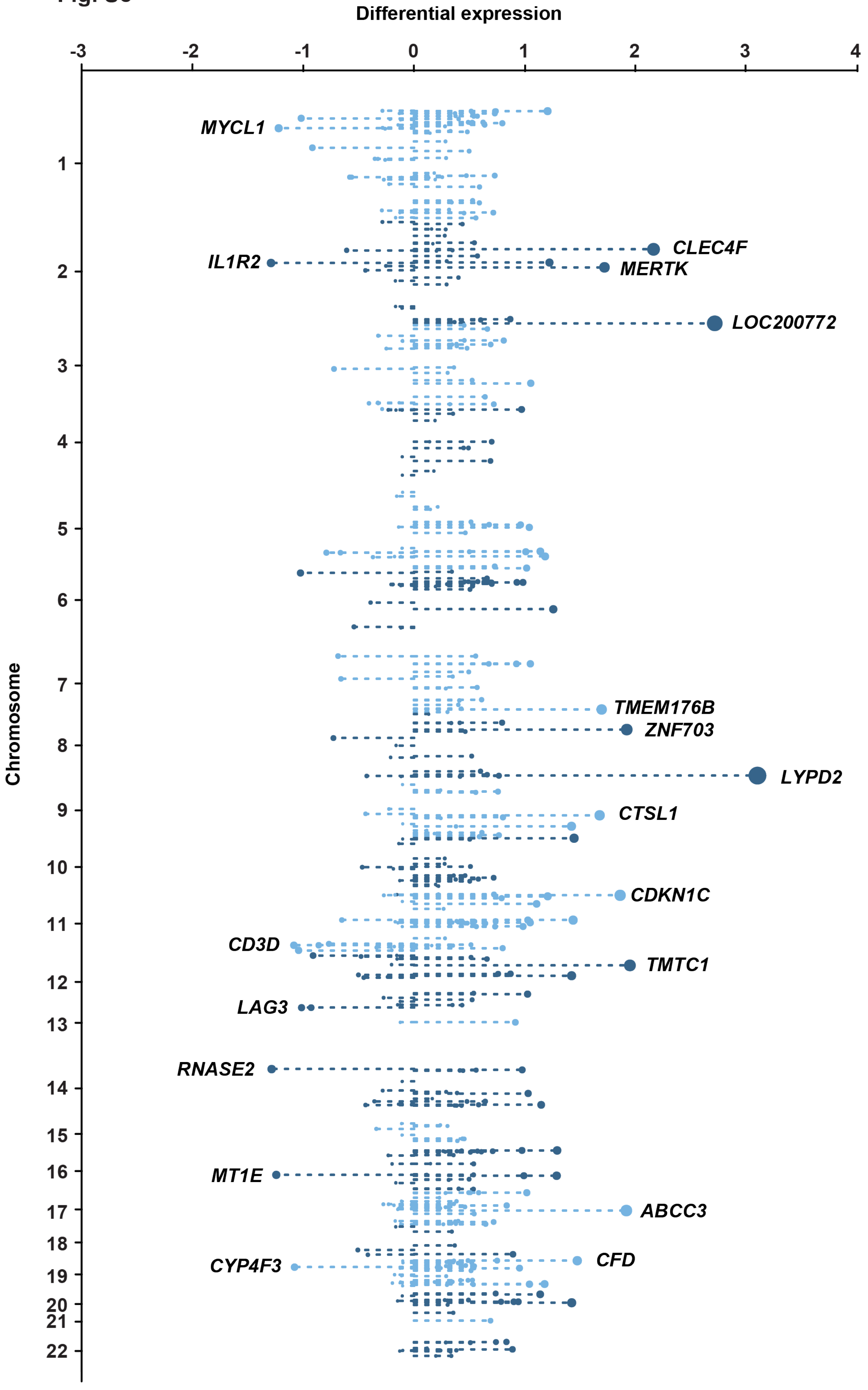

Differential expression
-3
-2
-1
0
1
2
3
4
Chromosome
MYCL1
IL1R2
CLEC4F
MERTK
LOC200772
TMEM176B
ZNF703
LYPD2
CTSL1
CDKN1C
CD3D
TMTC1
LAG3
RNASE2
MT1E
ABCC3
CFD
CYP4F3

**Supplementary Fig. 8**. Distribution of (autosomal) LATE genes plotted Manhattan-style along the genome, based on the largest differential expression observed across the measured cell types. Some of the most highly differentially expressed genes are located on chromosomes 2 (LOC200772, CLEC4F, MERTK, IL1R2) and 8 (LYPD2, ZNF703); however the figure also shows regions containing clusters of LATE genes, e.g. on chromosome 11.

### Fig. S9a

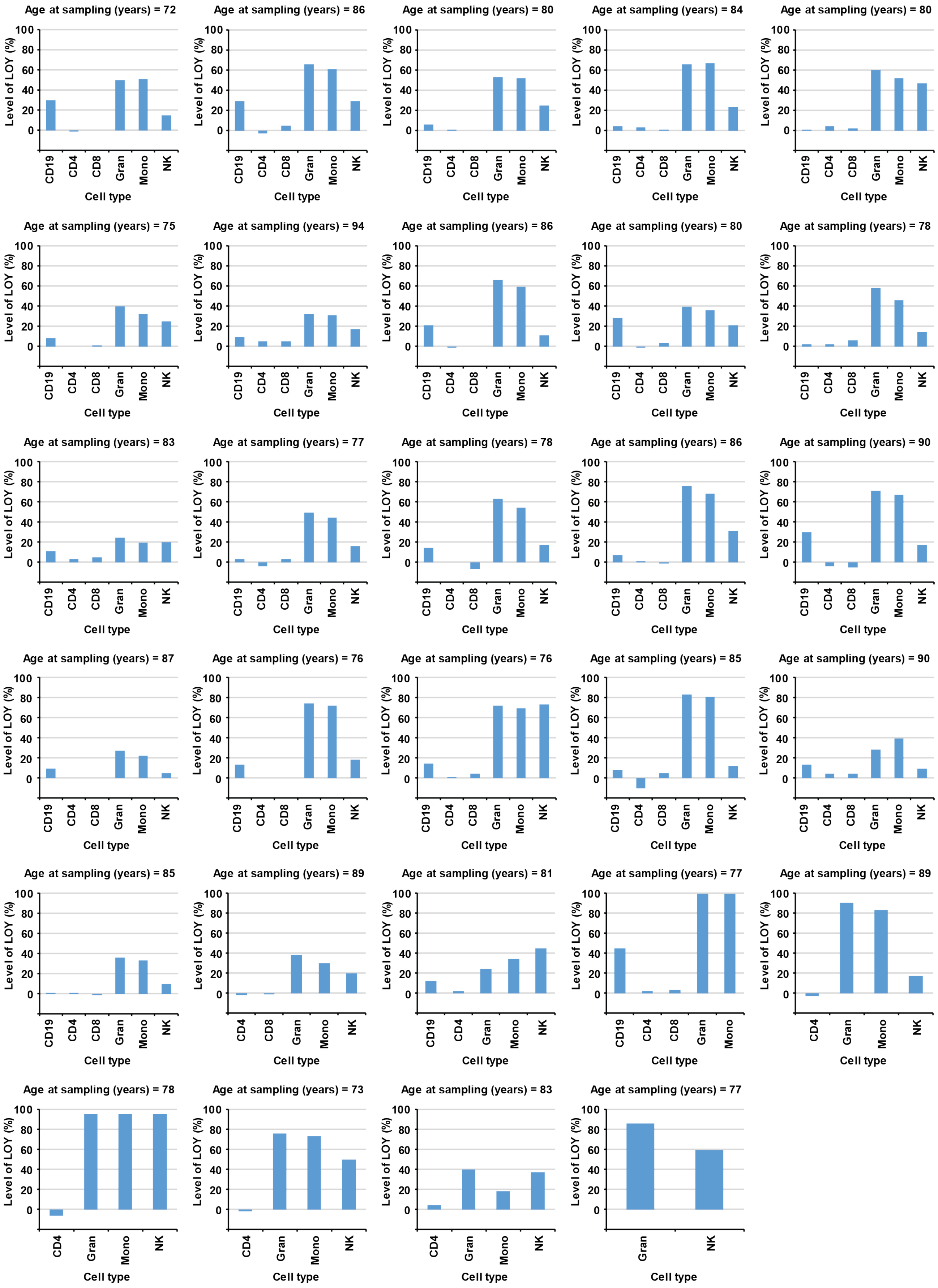

Age at sampling (years) = 72
Level of LOY (%)
100 80 60 40 20 0
CD19 CD4 CD8 Gran Mono NK
Cell type
Age at sampling (years) = 86
Level of LOY (%)
100 80 60 40 20 0
CD19 CD4 CD8 Gran Mono NK
Cell type
Age at sampling (years) = 80
Level of LOY (%)
100 80 60 40 20 0
CD19 CD4 CD8 Gran Mono NK
Cell type
Age at sampling (years) = 84
Level of LOY (%)
100 80 60 40 20 0
CD19 CD4 CD8 Gran Mono NK
Cell type
Age at sampling (years) = 80
Level of LOY (%)
100 80 60 40 20 0
CD19 CD4 CD8 Gran Mono NK
Cell type
Age at sampling (years) = 75
Level of LOY (%)
100 80 60 40 20 0
CD19 CD4 CD8 Gran Mono NK
Cell type
Age at sampling (years) = 94
Level of LOY (%)
100 80 60 40 20 0
CD19 CD4 CD8 Gran Mono NK
Cell type
Age at sampling (years) = 86
Level of LOY (%)
100 80 60 40 20 0
CD19 CD4 CD8 Gran Mono NK
Cell type
Age at sampling (years) = 80
Level of LOY (%)
100 80 60 40 20 0
CD19 CD4 CD8 Gran Mono NK
Cell type
Age at sampling (years) = 78
Level of LOY (%)
100 80 60 40 20 0
CD19 CD4 CD8 Gran Mono NK
Cell type
Age at sampling (years) = 83
Level of LOY (%)
100 80 60 40 20 0
CD19 CD4 CD8 Gran Mono NK
Cell type
Age at sampling (years) = 77
Level of LOY (%)
100 80 60 40 20 0
CD19 CD4 CD8 Gran Mono NK
Cell type
Age at sampling (years) = 78
Level of LOY (%)
100 80 60 40 20 0
CD19 CD4 CD8 Gran Mono NK
Cell type
Age at sampling (years) = 86
Level of LOY (%)
100 80 60 40 20 0
CD19 CD4 CD8 Gran Mono NK
Cell type
Age at sampling (years) = 90
Level of LOY (%)
100 80 60 40 20 0
CD19 CD4 CD8 Gran Mono NK
Cell type
Age at sampling (years) = 87
Level of LOY (%)
100 80 60 40 20 0
CD19 CD4 CD8 Gran Mono NK
Cell type
Age at sampling (years) = 76
Level of LOY (%)
100 80 60 40 20 0
CD19 CD4 CD8 Gran Mono NK
Cell type
Age at sampling (years) = 76
Level of LOY (%)
100 80 60 40 20 0
CD19 CD4 CD8 Gran Mono NK
Cell type
Age at sampling (years) = 85
Level of LOY (%)
100 80 60 40 20 0
CD19 CD4 CD8 Gran Mono NK
Cell type
Age at sampling (years) = 90
Level of LOY (%)
100 80 60 40 20 0
CD19 CD4 CD8 Gran Mono NK
Cell type
Age at sampling (years) = 85
Level of LOY (%)
100 80 60 40 20 0
CD19 CD4 CD8 Gran Mono NK
Cell type
Age at sampling (years) = 89
Level of LOY (%)
100 80 60 40 20 0
CD4 CD8 Gran Mono NK
Cell type
Age at sampling (years) = 81
Level of LOY (%)
100 80 60 40 20 0
CD19 CD4 Gran Mono NK
Cell type
Age at sampling (years) = 77
Level of LOY (%)
100 80 60 40 20 0
CD19 CD4 CD8 Gran Mono
Cell type
Age at sampling (years) = 89
Level of LOY (%)
100 80 60 40 20 0
CD4 Gran Mono NK
Cell type
Age at sampling (years) = 78
Level of LOY (%)
100 80 60 40 20 0
CD4 Gran Mono NK
Cell type
Age at sampling (years) = 73
Level of LOY (%)
100 80 60 40 20 0
CD4 Gran Mono NK
Cell type
Age at sampling (years) = 83
Level of LOY (%)
100 80 60 40 20 0
CD4 Gran Mono NK
Cell type
Age at sampling (years) = 77
Level of LOY (%)
100 80 60 40 20 0
Gran NK
Cell type

### Fig. S9b

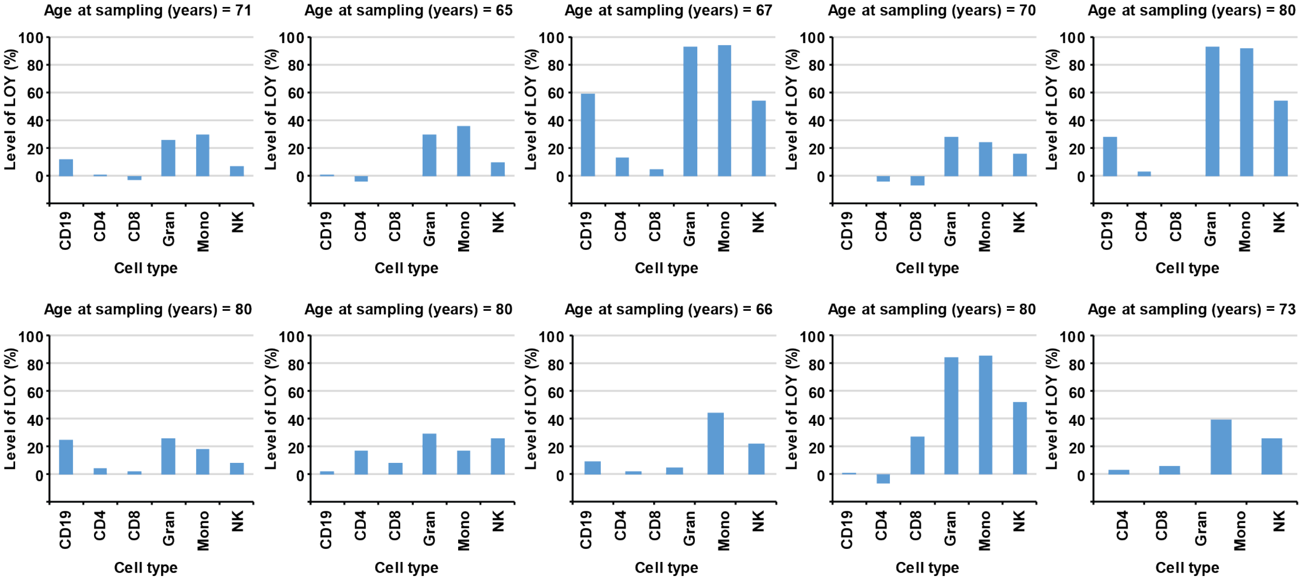

### Fig. S9c

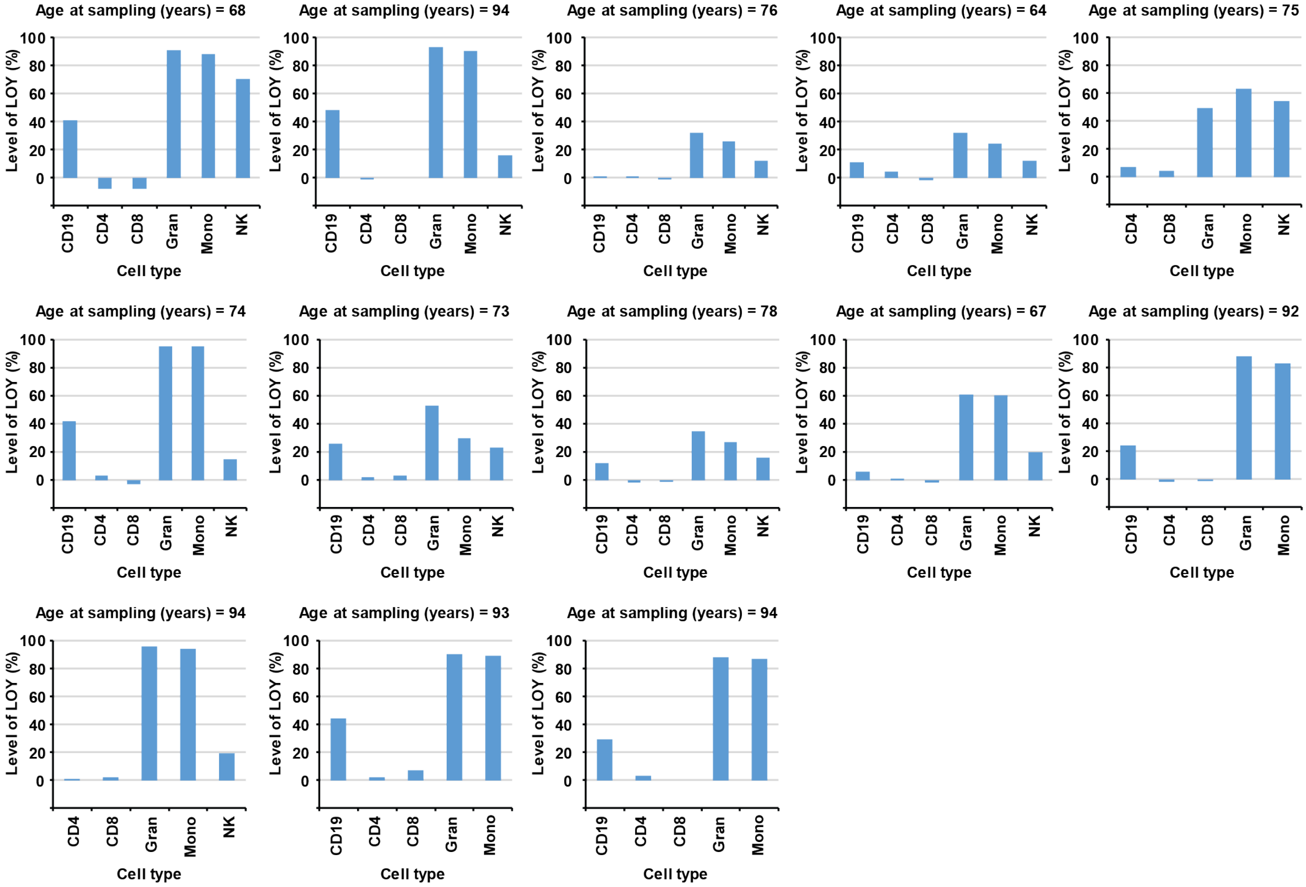

**Supplementary Fig. 9.** LOY oligoclonality observed in the AD and PC patients as well as controls. The level of LOY (y-axes) was estimated from DNA using SNP-arrays in up to six different types of sorted leukocytes per individual. LOY oligoclonality in each individual was defined as having LOY in >20% of cells in at least two types of studied leukocytes. **Panel a** show the level of LOY in different cell types observed in 29 men with LOY oligoclonality and diagnosed with Alzheimer's disease. **Panel b** display LOY in the studied cell types observed in 10 men with LOY oligoclonality and diagnosed with prostate cancer. **Panel c** show the level of LOY in 13 control subjects used in the analyses of Alzheimer's disease and prostate cancer. Abbreviations: AD = Alzheimer's disease, PC = prostate cancer, CD19 = CD19+ B-lymphocytes, CD4 = CD4+ T-lymphocytes, CD8 = CD8+ T-lymphocytes, Gran = granulocytes, Mono = monocytes and NK = natural killer cells.
